# A systematic approach to analyze T cell migration: application to mouse melanoma tumors

**DOI:** 10.1101/2025.04.29.651285

**Authors:** Nikolaos Memmos, Jason S. Mitchell, Brian T. Fife, David Masopust, David J. Odde

## Abstract

T cells must assess and choose between surveilling large areas, but also engage efficiently with the target cells. This process is translated into variations in speed and turning angle of T cells. In this study, we propose a generalized algorithm to analyze cell migration data with focus on CD8+ T cells, using clustering technique to identify the number of different migration states and Hidden Markov Model to capture the dynamical switching between them. The algorithm only requires a set of position observations in a series of times, independent of other factors. While this study focuses on CD8+ T cell migration, this approach can potentially be used broadly to study the migration of other cell types as well. For the current analysis, low and high avidity T cells in melanoma tumors were tracked *ex vivo* using two-photon microscopy. Our findings suggest that CD8+ T cells follow a two-state migration dynamic, with one state being faster, while the other slower and more localized. Moreover, we established a statistical methodology to analyze T cell migration to assess whether there is true variability in cell speeds as distinguished from stochastic fluctuations about a single speed, and it can be applied across different experimental platforms.

## Introduction

Cell migration is a crucial function across different living organisms and scales[1]. The effectiveness of the immune response depends on how efficiently specific types of immune cells, such as macrophages, Natural Killer (NK) cells, and CD8+ T cells, migrate to locate and eliminate pathogens in the body[2], [3], [4], [5], [6].

CD8+ T cells are part of the adaptive immune system and have a key role in immune surveillance for pathogens and tumor cells in the body. Their high specificity resulted in a rapid rise of CD8+ T cell-based cancer immunotherapies[7], [8], such as chimeric antigen T cell therapies and tumor-infiltrating lymphocyte therapies. The higher efficacy of T cell immunotherapies for hematological malignancies versus solid tumors, because of the easier tumor accessibility and better T cell survival, suggests the key role of CD8+ T cell migration as a limiting factor regarding the optimization of immunotherapies by tuning their ability to infiltrate and navigate into complex microenvironments for solid tumors[9], [10]. CD8+ T cells have been shown to be present in many different solid tumors, such as glioblastoma, melanoma, and lung squamous cell carcinoma[11], [12], [13]. Lymphocyte migration disruption, for instance, can be a potential target for anti-inflammatory drugs for diseases such as Crohn’s disease, multiple sclerosis, and asthma[14], [15], [16], [17], [18], [19]. Thus, understanding CD8+ T cell migration is crucial for optimizing the efficacy of immunotherapies and controlling the immune response generally.

Many statistical models have been proposed to describe cell motion during migration.

The simplest is the Brownian motion model, which was named after botanist Robert Brown who first observed that the motion of pollen into the water follows this pattern in 1827[20]. Later in 1855, Adolf Fick recognized that diffusion is analogous to heat transfer, so he proposed the diffusion equation to describe the transport of nutrients through membranes[21]. However, it was not until 1905 that Albert Einstein derived the diffusion equation in terms of probability density and postulated it[20], [22]. The three postulates are: i) two consecutive time intervals are independent and as a result uncorrelated, ii) displacements are identically distributed during each time interval and iii) the second moment of the displacements exist[23]. Brownian motion is characterized by a linear mean-squared displacement (MSD) with respect to time interval over which a displacement occurs. While this model is widely used in biology to describe the motion of cells, in reality, cell migration often violates the first postulate of the Brownian motion model and other models have been proposed, such as the persistent random walk (PRW) model, which parameterizes cell motility in terms of speed, *S*, and persistence time, *P*[24]. The PRW model is analogous to the Ornstein-Uhlenbeck stochastic process and has been used extensively to model cell migration when cells exhibit correlations in displacement, while the Brownian motion model is a subcase of the PRW model under the assumption that the system is overdamped[25] or when the sampling time interval is longer than the persistence time in the case of experimental data. It is characterized by a MSD proportional to the square of time interval at short times, and linear for long times. However, cell migration strongly depends on the properties of the extracellular environment and often deviates from the Brownian motion and PRW models. Wu *et al.* showed that cells can exhibit anisotropic behavior in 3D environments, compared to 2D environments[26]. Moreover, anomalous diffusion models have been proposed to explain experimental data that deviate from the linear relationship between MSD and time. Dieterich *et al.* studied the migration of NHE^+^ and NHE-deficient cells from the renal epithelial Madin– Darby canine kidney cell strains and concluded that both populations follow a superdiffusive migration pattern[27].

CD8+ T cells are a characteristic example where migration cannot be described by the traditional Brownian motion and PRW mathematical models. Boisfleury-Chevance *et al.* showed that lymphocytes exhibit a completely different migratory behavior compared to granulocytes and monocytes and speculated that lymphocytes follow a two-state migration pattern, while there is no closed-form mathematical model to describe their motility patterns, unless in the case of long times[28]. Sadjadi *et al.* also studied the migration of T cells in 3D collagen matrices and pointed out the existence of three motility types: slow, fast and mixed[29]. They proposed that slow moving T cells create channels for fast T cells in 3D matrices and their motion can be described by a two-state random walk. In both papers, however, the number of substates and the distributions of the observables (i.e. step length and turning angle) on each substate are not well-established.

Many different models have been proposed to explain CD8+ T cell migratory behavior.

For example, Beauchemin *et al.* proposed a model that is described by three parameters, the mean free time, *tfree*, a constant speed *vfree*, and a pause time *tpause*[30]. During the mean free time, the cell moves in a straight line with constant speed and pauses until it will pick a new random direction in which to move. However, the mean-squared displacement is sensitive to the number of data points, the signal-to-noise ratio, and the correlation between the MSD values, which can lead to incorrect estimations [31], [32], [33], [34]. A plethora of the proposed models have been initially used in animal ecology to explain the different motion patterns in animal movement.

Viswanathan *et al.* first proposed Lévy flight as model that can explain the searching process of animals for targets[35]. Lévy walk and Lévy flight are a class of random walk models where their step length, *λ*, distribution is characterized by a power-law relationship, *P*(*λ*)∼*λ*^−(1+*μ*)^, where μ is the power-low exponent and belongs to (0, 2][36]. These two terms are often used interchangeably in literature; however, they are distinct. Lévy flight assumes that the displacement between two points is discontinuous, compared to Lévy walk[37]. The argument for using Lévy walk over other models is that it optimizes the searching strategies for living organisms, when the targets (e.g., food) are sparse[38]. This model has also been used to explain the behavior of marine predators, bacteria, and human migration[39], [40], [41], [42]. Harris *et al.* studied the CD8+ T cell migration in mouse brains infected with the pathogen *toxoplasma gondii*[43]. They demonstrated that the chemokine CXCL10 enhances the ability of CD8+ T cells to efficiently identify the target cells, resulting in shorter capture times, and CD8+ T cells exhibit a generalized Lévy walk behavior. However, a lot of criticism arose about the Lévy walk model regarding both its biological and statistical grounding[44], [45], [46]. In particular, Lévy walk assumes that displacements are uncorrelated, while the step length, drawn from a heavy-tail distribution with infinite variance, can result in rare, unrealistically large displacements, even though this limitation can be overcome to some degree with truncated distributions. Furthermore, Edwards *et al.* also reported that the likelihood methods used to infer Lévy walk are inaccurate[45]. Banigan *et al.* proposed a Gaussian Mixture Model (GMM) to describe the non-Brownian behavior of CD8+ T cells in uninflamed lymph nodes[47]. While this model captures the existence of subpopulations based on their migration features, it fails to capture the possibility of the dynamic transitions between these populations. Jerison *et al.* showed that T cells in live larval zebrafish follow a two-state migration manner and proposed a variation of the PRW model, where persistence time, *P*, is linearly dependent on speed, *S*, which they called a speed-persistence coupling model[48]. While this speed-persistence coupling model is true in this system, it is not clear that it applies to other systems.

Hidden Markov Models (HMMs) have been widely used to describe the relationship between a sequence of observed data (observables) and events that are not observed directly (hidden events). HMMs were originally used in speech recognition but also gained popularity in studying biological sequences (e.g., DNA sequencing)[49], [50]. In cell migration, HMMs have been used to annotate and infer the dynamical switching between distinct morphological states, but also to describe the stochastic behavior of cell movement[51], [52], [53], [54]. For example, Degerman *et al.* applied a two-state HMM to glial progenitor cells and type-2 astrocytes and showed that the latter follow a directed movement pattern 2/3 of the time, compared to the first that behaved mostly as random walkers[51]. Torkashvand *et al.* proposed to initially cluster CD8+ T cells based on their motility pattern (Lévy and Brownian types) and the length of their step increments (short and long increments) to capture the heterogeneity across the cells[55].

This analysis resulted in 4 different clusters and inside each cluster homogeneous and non-homogeneous HMMs with 2 and 3 hidden states to capture the heterogeneity within the cell tracks. While in the current literature there is ample speculation of the existence of substates in T cell migration, there is no algorithm that rigorously establishes the minimum number of substates.

In this study, we propose a generalized algorithm to test for heterogeneity in cell migration and the possibility of dynamic transition between different speed states with application to CD8+ T cells *ex vivo* in a mouse melanoma model. We utilized published 3D two-photon microscopy data from melanoma tumors that include high and low avidity CD8+ T cells[56]. Avidity is defined as the total binding strength of the receptors of CD8+ T cells with the target cells. Our analysis shows CD8+ T cells, regardless of avidity type, move in both slow- and fast-moving states and switch dynamically between them.

## Materials and Methods

### T cell migration in mouse melanoma tumors data

Four datasets from B16F10 melanoma tumors from transgenic mice were obtained via *ex vivo* two-photon microscopy, which were previously published by Tucker *et al.*[56]. The transgenic mice produced both low and high avidity CD8+ T cells which were specifically recognizing the tumor Ag tyrosinase-related protein 2 (TRP2). The CD8+ T cells were purified from the spleen and lymph node and then stimulated in vitro with interleukin-12 (IL-12) to prevent exhaustion. Subsequently, the CD8+ T cells were transferred intravenously to the mice. The low and high avidity CD8+ T cells were identified with Brainbow tdTomato and UBC-GFP respectively and tracked in 3D with Imaris software with a frame rate equal to 32 seconds/frame. To reduce bias, only tracks with consecutive time points were allowed in the analysis, otherwise they were considered as separate tracks (e.g. so there were no missing points within one track). In total, N=271 low and N=266 high avidity CD8+ T cells, with 5204 and 5067 total time points respectively, were analyzed. Because there was potentially a significant difference between the migration pattern of the two populations, low and high avidity CD8+ T cell populations were analyzed separately. The total imaging time and time window were similar for the high- and low-avidity datasets (**Table S1**).

### Statistical analysis

The time averaged mean-squared displacement (TAMSD) and the displacement autocorrelation function were calculated for each individual cell. Assuming a time series of 3D positions *r*_1_, *r*_2_, …, *r*_*N*_ with N data points *k* equally spaced time lags, where each vector ***ri***, corresponds to the position of the cell, ***ri*** *= (xi, yi, zi)*, at time point *i*, the MSD was estimated using the overlapping sampling method[57]:

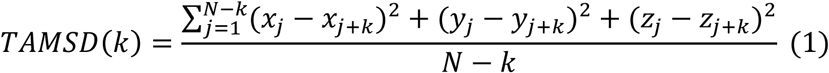

Displacements were calculated with first differencing from the position datasets to obtain detrended time series. Thus, it was assumed that the time series of the displacements have a mean equal to zero, which is valid for detrended datasets[58], where the autocorrelation function for each individual cell is given by:

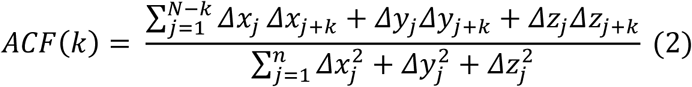

To assess autocorrelation function at the population level, the weighted average ACF based on the number of data points for each cell was estimated. The 95% confidence interval was calculated based on Equation S1.

### Gaussian Mixture Model

The Gaussian Mixture Model (GMM) was applied to the distribution of the displacements of the total number of CD8+ T cells and all time points to identify the number of migration states[47]. In this clustering method, the distribution is approximated as a linear superposition of *M* Gaussian distributions:

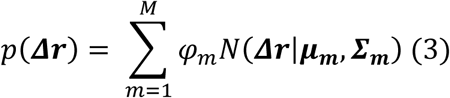

where ***Δr*** are the displacements, *φk* are the mixing coefficients and must obey the following conditions:

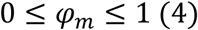

and

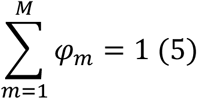

For the purposes of this study, it was assumed that the mean of the displacement Gaussians, *μk*, is zero and motion is isotropic, so that equation 3 can be written as:

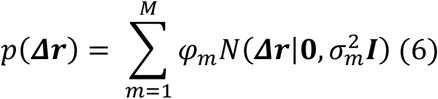

To identify the set of parameters {*φm*, σ^2^_*m*_ } which maximize the likelihood of the distribution of the displacements (equation 7), the Expectation-Maximization algorithm has been used[59].

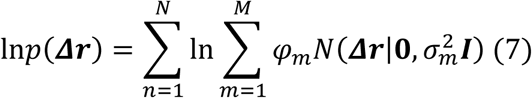

The Bayesian Information Criterion (BIC) and Akaike Information Criterion (AIC) along with the Elbow Method were used to decide the number of states[60].

### Hidden Markov Model

To infer the parameters of possible dynamic switching between states, a Hidden Markov Model (HMM) was applied, using the *momentuHMM* package, in combination with modified scripts from the *moveHMM* package, in R language[61], [62]. The number of hidden states was dictated by the clustering analysis with GMM as described previously[47]. The observable variables of the model are {*s, θ, Δz*}, where s is the step length of a cell projected into the xy-plane, *θ* is the turning angle projected onto the xy plane and *Δz* is the displacement in the direction of the z-axis. To fit the HMM, it was assumed that *s*, *θ* and *Δz* follow a gamma distribution, von Mises distribution, and logistic distribution, respectively:

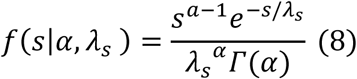

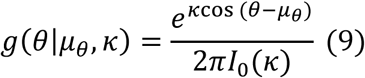

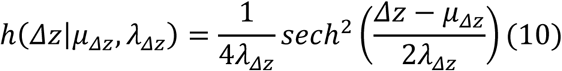

where *α and λs* are the shape and scale of Gamma distribution respectively, *μθ* and *κ* are the location and the concentration parameters of the Von Mises distribution, and *μΔz* and *λΔz* are the location and scale parameters of the logistic distribution, respectively. The caveat of the gamma distribution is that it cannot model steps with zero length, thus a zero-inflated distribution is needed, which assigns a probability *z* to observe a zero value and a *1-z* probability to observe a positive value based on a gamma distribution[61], [63]. It was assumed that the von Mises distribution is symmetric around zero, so *μ*_*θ*_ = 0. The turning angle, θ, was calculated using the *atan2* function to constrain it in the (-*π*, *π*] range:

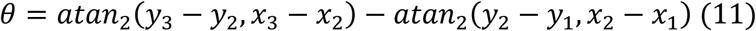

where *x, y* are the xy-coordinates, indices 1, 2, and 3 correspond to 3 consecutive time points in a cell track. To estimate the parameters, a numerical maximization of the likelihood was employed[64], [65].

## Results

### T cell migration heterogeneity

Infiltrated CD8+ T cells in B16 melanoma tumors were tracked using ImageJ for a period of 30 minutes at a frame time interval of 32 seconds. The red cells correspond to low avidity T cells, while the green ones correspond to high avidity T cells (**Figure 1A**). Compared to other types of immune cells, like monocytes and granulocytes, both T cell populations follow a distinctive migration pattern that does not appear to be described by a single migration speed, but instead appears to migrate in a bimodal pattern, consisting of distinct slow- and fast-moving states (**Figure 1B-C**)[28]. The slow-moving state tends to be more localized, possibly because of the engagement of T cells with target cells (e.g. dendritic and tumor cells). On the other hand, the fast-moving state is persistent in a certain direction.

**Figure 1.**
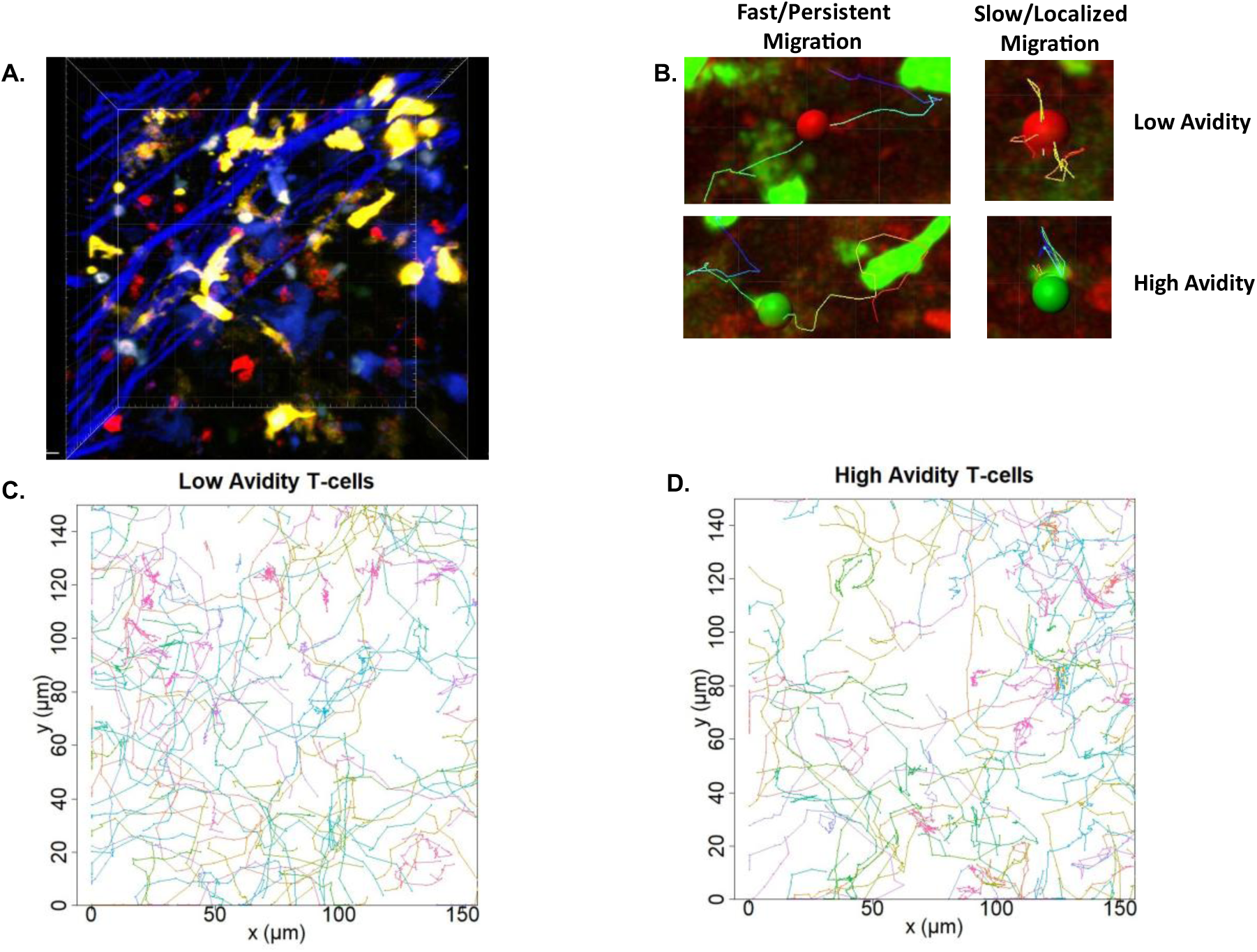
**A**) Example of a mouse melanoma migration dataset where green cells correspond to high avidity CD8+ T cells, red to low avidity CD8+ T cells, blue channel on collagen and yellow on dendritic cells. **B**) Examples of fast/persistent (left) and slow/localized (right) migration patterns are evident for both low and high avidity T cells. **C-D**) An xy-projection of (**C**) low and (**D**) high avidity CD8+ T cell migratory tracks. It appears by eye that the cells migrate in either a localized (slow) or persistent (fast) manner. .

The distribution of the displacements of the low and high avidity CD8+ T cells across *x*,*y* and *z* were compared using the Kolmogorov-Smirnov (KS) test to decide if the two populations should be analyzed together. The KS test revealed that the two populations differ, with p-values equal to 0.024, 0.0001 and 0.016 for *x*, *y* and *z* axes respectively, which may be attributed to the different dissociation rates (**Figure S1**).

To assess whether there is variability with respect to migration pattern among individual T cells, the time averaged mean-squared displacement for each individual cell (TAMSD, **Figure 2A**) was calculated based on equation (1). The average displacement autocorrelation function for the T cell populations was also estimated, based on equation (2), to evaluate whether or not T cells exhibit directional memory (ACF, **Figure 2B-C**). The TAMSD is a first estimator of the variability in T cell motility behavior, while the average autocorrelation function describes the directionality and persistence of the T cell migration. **Figure 2A** shows that TAMSD can vary largely between CD8+ T cells, where low values suggest that T cells perform a localized searching because of their interactions with target cells, while increasing values correspond to a persistent random walk. However, such variations on a cell-to-cell basis can also result from a single random walk model applying to all cells due to stochastic fluctuations. On a population level, the average autocorrelation function was calculated and fitted to a simple exponential function, *e*^−*t*/*τ*^, where *τ* is the decay time constant. For the low avidity T cells, the decay time constant is *τ*_*low*_ = 0.690 ± 0.035 minutes and for the high avidity, *τ*_*high*_ = 0.632 ± 0.030 minutes, again confirming that migration differences between high- and low-affinity T cells are modest. The low avidity T cells exhibit a slightly higher decay time constant, most likely because they spend more time in a persistent migration pattern without efficiently associating with target cells. Both numbers are close to what has been previously observed experimentally for lymphocyte migration *in vitro*[28].

**Figure 2.**
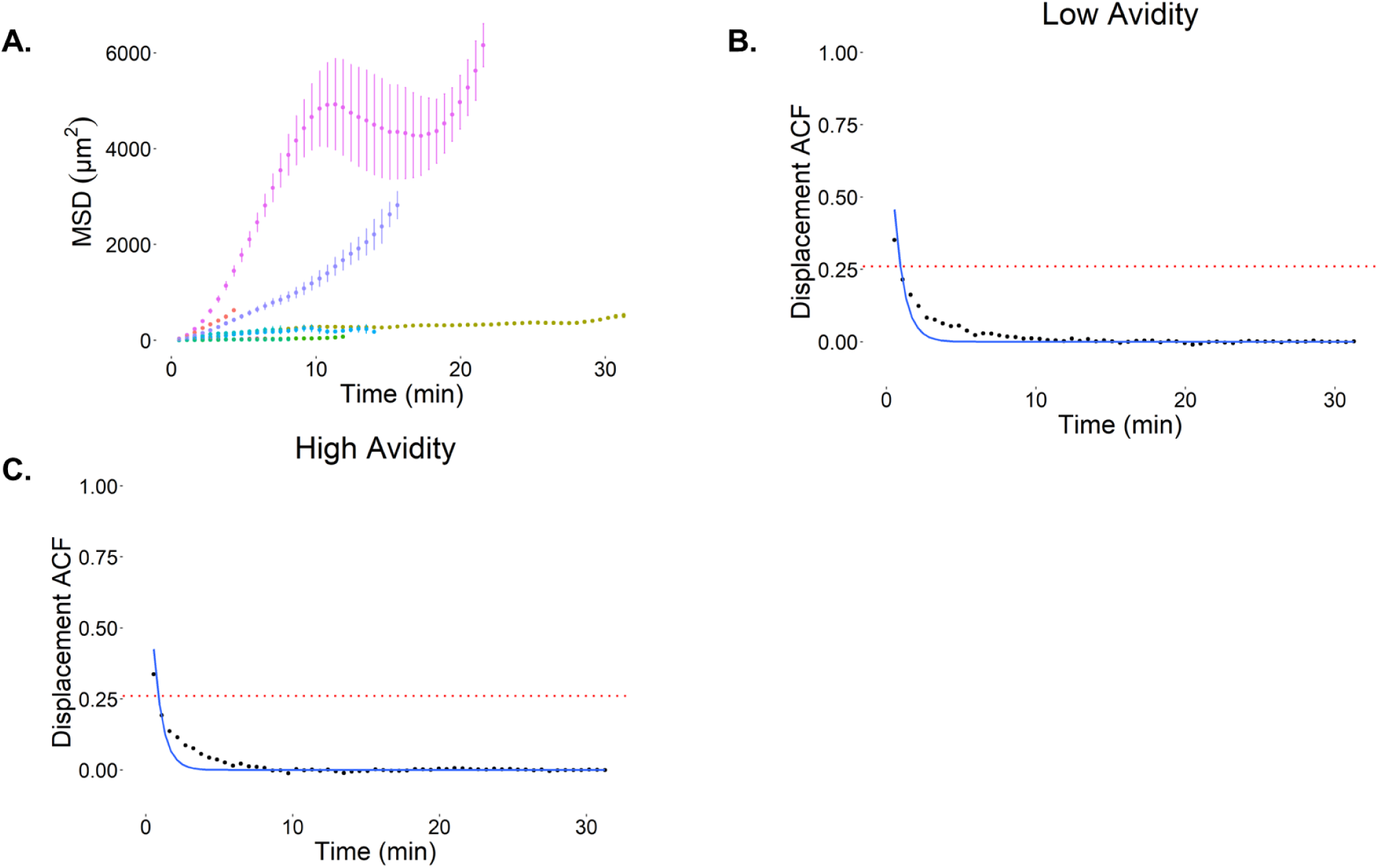
(**A**) Selected examples of the time averaged mean-squared displacement (TAMSD) of individual CD8+ T cells and (**B-C**) the average autocorrelation function over all high and low avidity T cells respectively. The average time window between the time points is 32.339 sec. The black points are the experimental data, the blue curve the fitted exponential function, *e*^−*t*/*τ*^, and the red dashed line corresponding to the 95% confidence interval. The *τ*_*low*_is equal to 0.6901 ± 0.0348 minutes and the *τ*_*high*_ to 0.6319 ± 0.0301 minutes.

### The normality tests reject the possibility of a single Gaussian

In the case of Brownian motion, the distribution of displacements follows a Gaussian distribution:

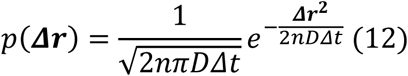

where ***Δr*** is the displacement of T cells, *n* is the number of dimensions, *D* is the diffusion (or random motility) coefficient, and *Δt* is the time increment. To evaluate if the T cells obeyed a Brownian motion pattern at the population level, we performed normality tests based on the total distribution of the displacements (**Figure 3A, C**) for each axis, using a QQ-plot (**Figure 3B, D**) and Anderson-Darling test (**Table S3**). Based on the QQ-plots, it is shown that the data (red points) diverge significantly from the Gaussian distribution (black line) across every direction and for both low and high avidity T cells. The deviation in both the positive and negative tails of the distribution suggests that the distributions are heavy tailed relative to a Gaussian. The same conclusion is drawn from the Anderson-Darling normality test, which rejects the null hypothesis of normality. Overall, we conclude that T cell migration is not well described by a random walk with q single random motility coefficient.

**Figure 3.**
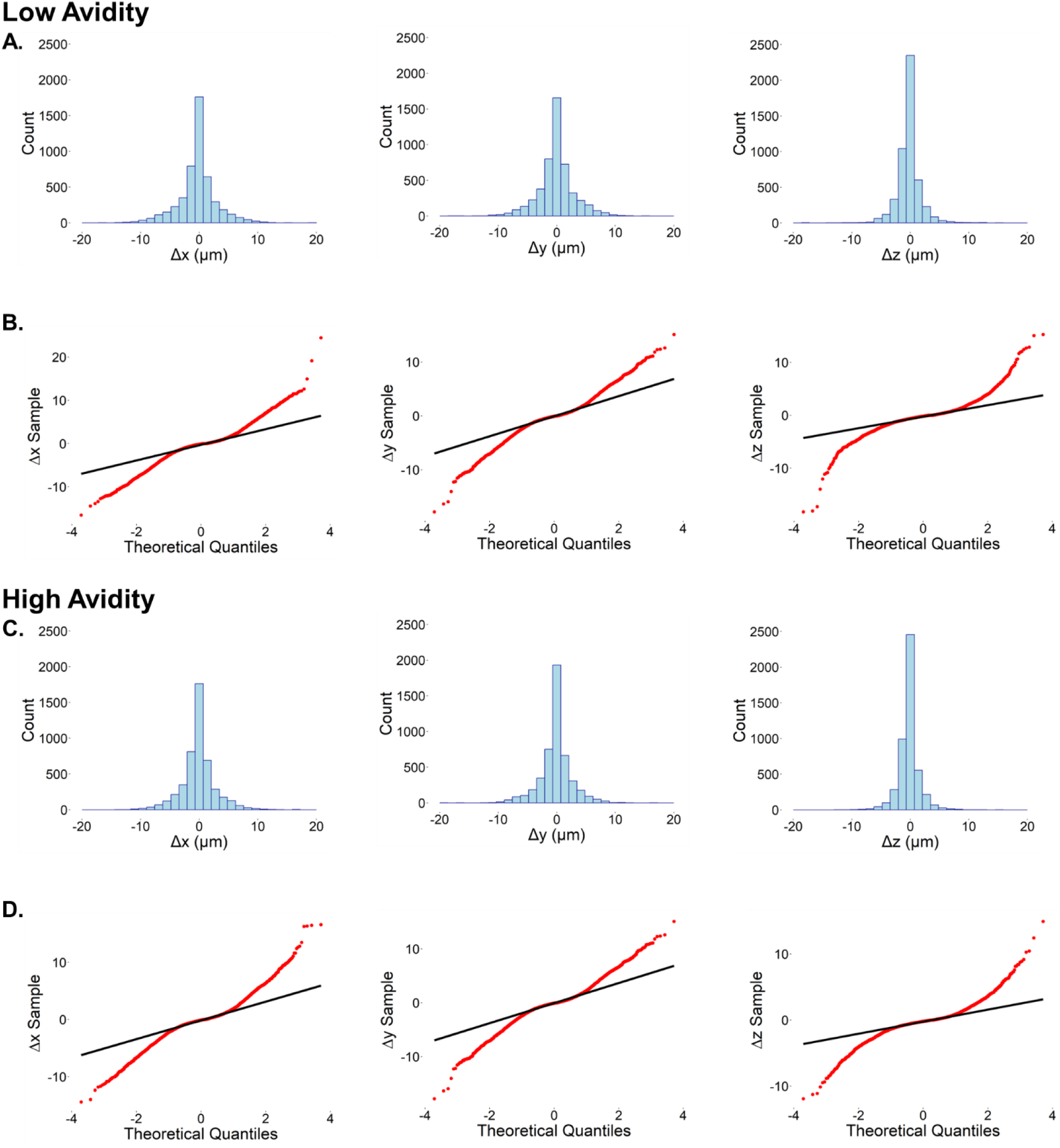
(**A, C**) The distribution of the displacements for low avidity CD8+ T cells and all time points across each axis for low and high avidity T cells respectively. (**B, D**) The corresponding QQ-plots for each distribution. The black line corresponds to theoretical expectation for a normal distribution and red points correspond to the data. The deviations at the distribution extremes implies that displacements are not well-described by a Gaussian distribution, but rather are more heavy-tailed.

### The Gaussian Mixture Model indicates a two-state migration

Since the normality tests rejected the possibility of a universal migration pattern described as Brownian motion, the question remains if a multiple state model can be used to model the migration of T cells. The existence of a heavy tailed displacement distribution suggests that T cell migration can be described by a Lévy walk[43]. However, this model has been criticized for its ability to describe foraging processes, because of the infinite variances and the inappropriate use of likelihood function [44], [45], [46]. Previously, Banigan *et al*. used a Gaussian Mixture Model to describe T cell migration in the lymph node in the absence of inflammation[47]. In our case, we used the same approach to reveal how many clusters of migration states exist in both T cell populations, but may not be a completely adequate model, since it does not capture the potential for transition between the faster and slower states. The relatively weak correlation in the displacements (**Figure 2B-C**), shows the weak persistence in the population migration behavior and justifies the use of GMM as a clustering method. To perform the clustering, it was assumed that the mean of the distributions is equal to zero and the only free parameters were the weights and the covariance matrices for each component.

Moreover, the assumption of isotropic migration of the CD8+ T cells in the tumors results in less free parameters, since the covariance can be described with a single parameter and takes the form *σ*^2^*I*, where *I* is the identity matrix. The tolerance for convergence of the maximum likelihood was set to 10^-6^ (**Figure S2**).

Both Akaike and Bayesian information criteria indicated that the optimal number of clusters to describe the probabilities was equal to two based on the elbow method (**Figure 4**). Higher numbers of states do not reduce the AIC and BIC further, indicating that two speeds (one fast, one slow) is the minimum number of substates. Our result of two substates agrees with Banigan *et al*., who also identified two T cell speeds[47]. From **Tables 1** and **2**, it is shown that a similar percentage of low and high avidity T cells belong to each state, however low avidity T cells appear to have larger variances on each state, because of their slightly higher speed, compared to high avidity T cells. While helpful in testing whether the data can be explained by a single migration model or whether substates exist, GMM does not capture potential dynamic switching between the states and should be used only as a clustering method when appropriate to determine the number of distinct migration speeds justified by the data.

**Figure 4.**
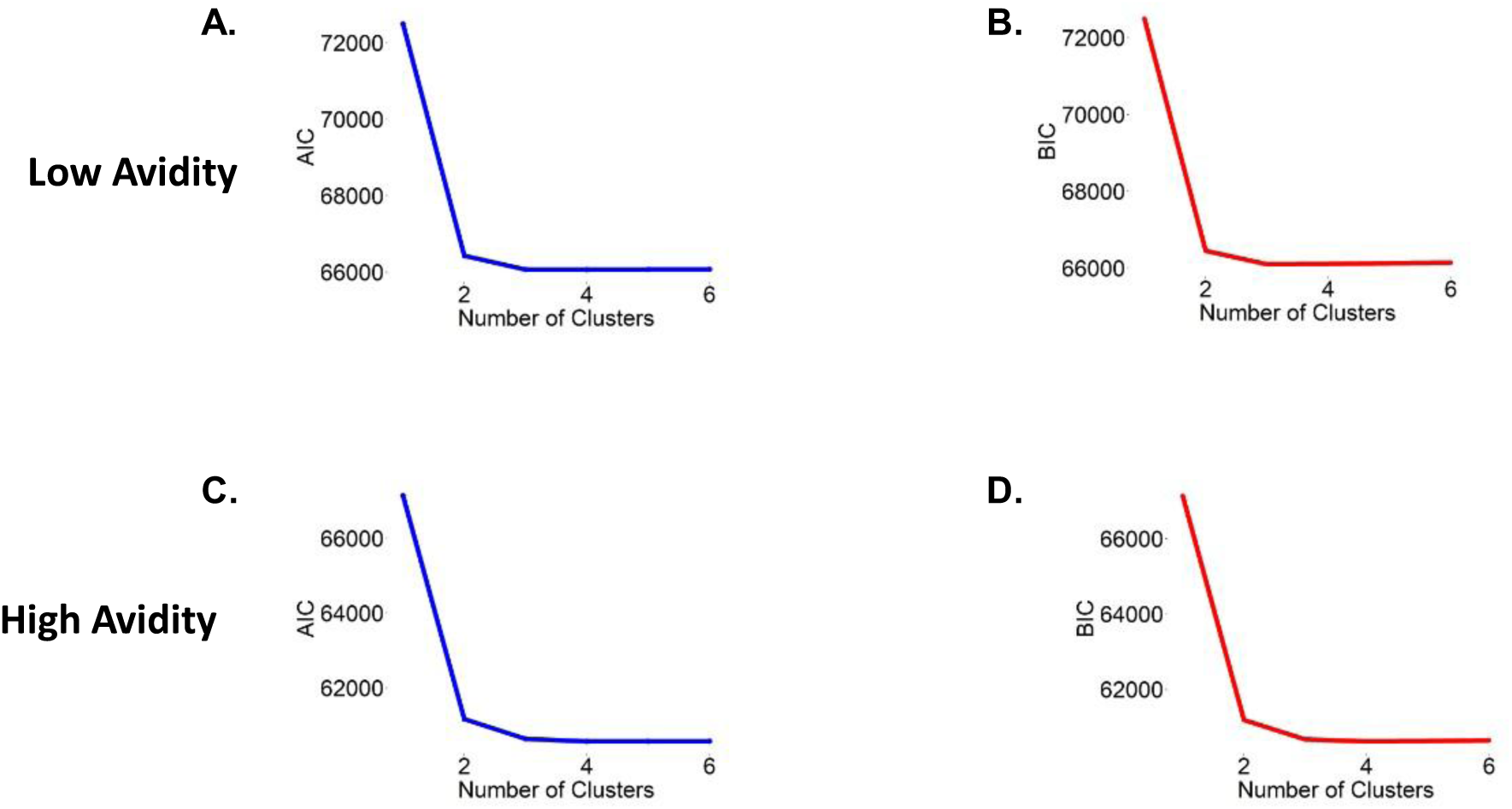
(**A-B**) The AIC and BIC scores with respect to number of components of the Gaussian Mixture Model for low and (**C-D**) high avidity CD8+ T cells. Both criteria indicate that the distribution of the displacements is better described with a two-state model, based on the elbow method.

### The Hidden Markov Model captures the dynamic behavior of T cells

The GMM indicated the existence of a two-state migratory behavior of T cells (i.e. a fast state and a slow state), however it cannot capture their dynamic behavior. For that reason, a Hidden Markov Model was applied to identify the transition probabilities between the fast and slow states. The observables for the HMM were the step length of the T cells in the *xy*-plane, the turning angle and the displacement in the *z*-axis. The probability distributions of these observables are described by the equations (8)-(10). In **Figure 5** it is shown that the step length distribution in the fast-moving state is broader (i.e. is faster, as expected), while the angle distribution is concentrated around zero (i.e. is more persistent, tending not to turn as much).

**Figure 5.**
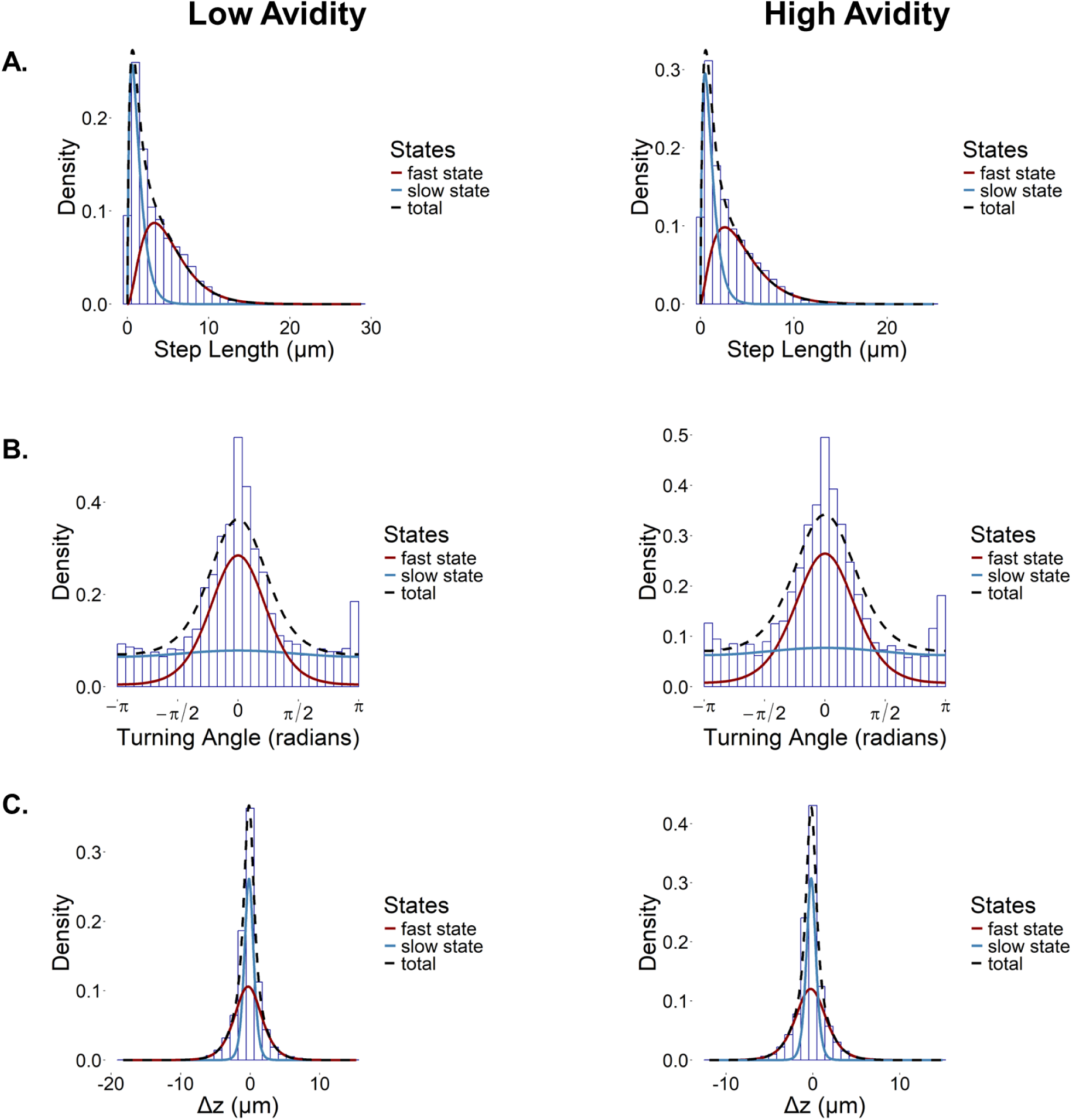
(**A**) The step distribution, (**B**) the turning angle distribution and (**C**) the distribution of the displacements across the z-axis of high avidity T cells. The red and blue lines correspond to the fast- and slow-moving states respectively. The dark red curves correspond to the slow-moving state, the blue to the fast-moving state and the black to the total density.

This observation confirms our initial speculation, that persistent motion correlates with faster migration. On the other hand, slow migration is accompanied by an almost uniform turning angle distribution, consistent with a random exploration of a small region in space. Furthermore, low avidity T cells move somewhat faster than high avidity T cells, in both their fast state and their slow state (Supplementary Information). **Tables 3** and **4** show the transition probabilities between the states for low and high avidity T cells, respectively. Each row corresponds to the current state and each column to the potential next state, while the sum of each row’s elements is equal to one (i.e. they must either remain in their current state or switch to the other state). We find that the average probability for a T cell to change state is equal to 0.136, which can be translated to a transition rate of 0.25 min^-1^ approximately (or equivalently switching every ∼4 min). Thus, the probability of T cells changing state, in both directions, in one minute is around 25%, which is independent of their avidity.

The fitted models were evaluated using their residuals (Supplementary **Figure S3-S4**). The QQ-plots of the pseudo-residuals, also known as quantile residuals, show that they are in good agreement with a normal distribution, while the ACF of the pseudo-residuals is weak for every observable, suggesting a good fit of the HMM to the data. In the case of high avidity T cells, the ACF of the pseudo-residuals for the step length, there is a relatively higher autocorrelation which can be attributed to their stronger interactions with the target cells or other environmental variables. However, in both cases, we consider that the HMMs satisfy the goodness-of-fit test, using the pseudo-residuals[65].

### The Hidden Markov Models capture the behavior of the original dataset

To validate the fitted Hidden Markov Model for each T cell population, a simulation of 100 T cells for 500 time points, using the fitted HMMs, was performed and the average ACF function was calculated (**Figure 6**). The time window for the simulations was assumed to be equal to the experimental dataset. Then a single exponential function was fitted as in the experimental data (**Figure 3**). As it is shown in **Figure 6**, the exponential function, *e*^−*t*/*τ*^, perfectly fits the simulated data. For the low avidity simulated T cells, the time decay constant is 0.701 ± 0.008 minutes and 0.587 ± 0.006 for the high avidity cells. In the case of low avidity T cells, the fitted time decay constant of the simulated data is closer to experimental one (*τ*_*low*_ = 0.6901 minutes), compared to the high avidity T cells (*τ*_*high*_ = 0.632 minutes). This discrepancy is a consequence of the higher autocorrelation in the step length for the high avidity model (**Figure S4**). In both cases, the average ACFs of the simulated data agree with the experimental ones. Moreover, the fitted HMMs are capturing the same behavior as in the experimental data, with respect to time averaged mean-squared displacement (**Figure S5**).

**Figure 6.**
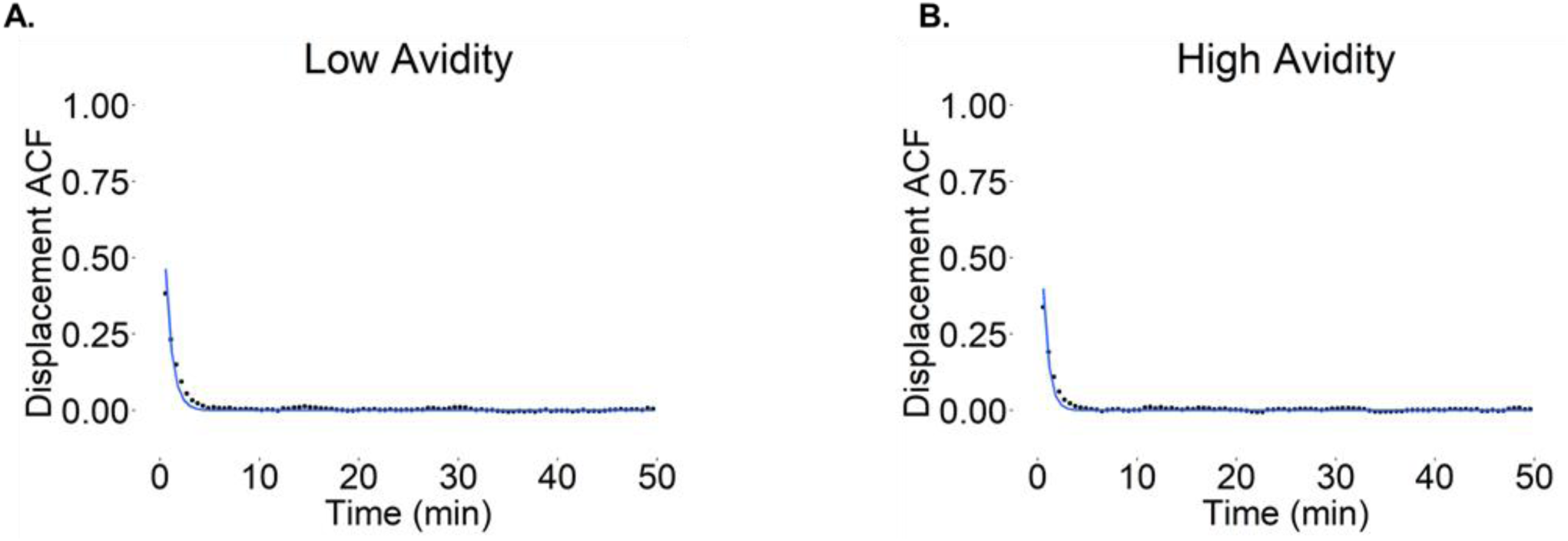
Autocorrelation Function of displacements shows only weak and short time-scale positive correlations. The fitted ACF on the simulation data for (A) low and (B) high avidity CD8+ T cells. The blue line corresponds to the fitted curve, the black points on the calculated points and the red line on the significance threshold. For each simulation, 100 cells were used and 500 time points. The time window was assumed to be equal to 32.339 seconds, as in the experimental dataset.

## Discussion

CD8+ T cells play a key role in surveilling the human body and eliminating pathogens and cancer cells. The rapid development of T cell-based cancer immunotherapies dictates the necessity for better engineering of CD8+ T cells to efficiently explore the tumor microenvironment and kill cancer cells. This can be achieved by properly tuning the migration of T cells in tumor microenvironments. Thus, the understanding of the nature of the movement of T cells, the existence of possible substates and their statistical properties is crucial for cancer immunotherapy optimization.

In this study, two T cell populations with different avidities were analyzed with respect to their migratory behavior in B16 melanoma tumors, using *ex vivo* 3D two-photon microscopy imaging [56]. Both T cell populations displayed two-state migration, with one state being slow and resembling a random walk, while the other being fast and persistent.

While these findings have also been observed in previous studies, here we propose a systematic algorithm and approach for analyzing T cell migration data to rigorously identify the number of migration substates, their properties, and the rates of transition among them. Because of the complexity of T cell migration and their importance, numerous mathematical models have been proposed to describe their behavior. It has been shown that naïve T cells follow a diffusive or subdiffusive random walk across different studies[66], [67], [68], [69]. On the other hand, when T cells are activated, they switch to more complex migration dynamics, which includes both ballistic-type fast migration (with long-tailed displacement distributions) and more localized (slow) search [69], [70]. This observation led to the use of the Lévy walk model to describe T cell migration in brain tissue, in response to infection[43]. While this model seems to explain the behavior of T cells, it fails to capture the autocorrelation between the displacements, which contradicts what happens in nature and based on our analysis, and it has been criticized based on the inappropriate use of likelihood function [45], [46], [71]. Edwards *et al*. demonstrated that previous studies used the residuals of the regression to estimate the likelihood function of the proposed models, instead of the likelihood of the actual probability distribution, which lead to misleading information criteria[45]. Furthermore, its validity has not been verified in other tissues[69]. Banigan *et al*. has shown that T cells do not follow a Lévy walk in a brain infection model, but a combination of slow- and fast-moving states, using a Gaussian Mixture Model, while Jerison *et al.* used the speed-persistence coupling model to describe this behavior[47], [48]. The findings of these papers agree with what we observe in our case with melanoma-infiltrating CD8+ T cells, but their model is not able to capture the switching between the states, which we now capture with the Hidden Markov Modeling approach. Moreover, the Hidden Markov Model can be applied across different conditions and platforms, compared to a closed-form equation. The similarity of the fast-slow substates suggests that T cell migration mechanics are conserved across infections and tumors.

In our analysis, we combined GMM and HMM, to analyze T cell migration in solid tumors, while trying to simultaneously minimize the number of assumptions. Our analysis indicated that CD8+ T cells, in both cases, do not obey pure Brownian motion based on normality tests, while the coexistence of two states was verified with a Gaussian Mixture Model, as in Banigan *et al*.[47]. The use of this clustering methodology is justified by the weak displacement autocorrelations. It was observed that low avidity T cells have a longer memory, compared to the high avidity T cells, potentially because of their weaker engagement with the target cells. The clustering analysis identified the number of hidden states that was used in the Hidden Markov Model to model the dynamic switching between the states. Finally, the fitted models were judged based on their ability to capture the behavior of the original dataset, using as metrics primarily the ACF and secondly, the reproduction of the TAMSD curves.

The understanding of CD8+ T cell migration can assist in the design of better immunotherapies for cancer treatment. Our analysis rigorously showed that T cells migrate in a two-state model, a slow-moving state with less directional persistence and a fast-moving state with high persistence, and that single cells can switch between these states. T cells can be engineered accordingly to infiltrate the tumor microenvironment and engage with the target cells, which surveil the area effectively. The exact nature is likely dependent on the type of tumor, for example solid tumors may require T cells to remain more in the slow-moving state after infiltration, because of the high density of target cells.

Even though the algorithm was applied to CD8+ T cells, the same approach can be applied to other cell types. The two major caveats of our algorithm are the use of the GMM which can only be applied in the case with low autocorrelation with regards to the displacement of the cells. If higher autocorrelations exist, other clustering methods, such as Dynamic Time Wrapping (DTW) should be used to identify the number of the states[72]. The second caveat is that it does not account for the influence of external factors, such as the effect of other types of cells, and the spatial component. Overall, our method can capture the features of T cell migration in tumor tissue *ex vivo*, without relying on closed-form equation models and with a small number of assumptions.

## Data availability statement

The data that support the findings of this study are available from the corresponding author upon reasonable request.

## Ethics approval statement

Experiments were conducted under specific pathogen-free conditions and performed in compliance with relevant laws and guidelines, with approval of the Institutional Animal Care and Use Committee at the University of Minnesota.

**Table 1.** The weights (*φ*) and the variances (*σ^2^*) of the two Gaussian mixture components for low avidity T cells.

| Low avidity T<br>cells parameters | Fast moving | Slow moving |
| --- | --- | --- |
| $\varphi$ | 0.562 | 0.439 |
| $\sigma^2 (\mu m^2)$ | 13.407 | 0.695 |

**Table 2.** The weights (φ) and the variances (σ^2^) of the two Gaussian mixture components for high avidity T cells.

| High avidity T<br>cells parameters | Fast moving | Slow moving |
| --- | --- | --- |
| $\varphi$ | 0.571 | 0.429 |
| $\sigma^2 (\mu m^2)$ | 10.450 | 0.512 |

**Table 3.** The transition rates of low avidity T cells given in min^-1^, based on the *r* = *p*/*Δt* equation, where *r* is the transition rate, *p* the probability, and *Δt* the experimental time window equal to 32.339 sec. The numbers within the parenthesis correspond to the probability values.

|  | Fast moving | Slow moving |
| --- | --- | --- |
| Fast moving | 1.596 $\text{min}^{-1}$<br>(0.860) | 0.258 $\text{min}^{-1}$<br>(0.140) |
| Slow moving | 0.222 $\text{min}^{-1}$<br>(0.125) | 1.626 $\text{min}^{-1}$<br>(0.875) |

**Table 4.** The transition rates of high avidity T cells given in min^-1^, based on the *r* = *p*/*Δt* equation, where *r* is the transition rate, *p* the probability, and *Δt* the experimental time window equal to 32.339 sec. The numbers within the parenthesis correspond to the probability values.

|  | Fast moving | Slow moving |
| --- | --- | --- |
| Fast moving | 1.584 $\text{min}^{-1}$<br>(0.853) | 0.270 $\text{min}^{-1}$<br>(0.147) |
| Slow moving | 0.240 $\text{min}^{-1}$<br>(0.130) | 1.614 $\text{min}^{-1}$<br>(0.870) |

## Acknowledgements

This work was supported by grants from the National Institutes of Health (grant numbers R01 AI156276 to B.T.F., U54CA210190, U54CA268069, and P01CA254849 to D.J.O.). Multiphoton imaging on the Leica Stellaris 8 DIVE system was made possible by a NIH S10 grant 1S10OD034456-01 (B.T.F.) to the University of Minnesota Center for Immunology.

## Table of Contents

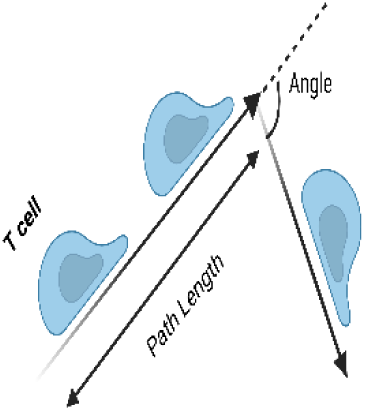

The results show that a minimum of two migration speeds can be rigorously identified from the data with cells switching between a fast, persistent migratory state, and a slow, random migration state. These results will help in identifying genetic factors that influence rapid migration, among other applications, such as quality control for CAR-T cell migration.

## Supplementary Information

To calculate the 95% confidence interval for white noise autocorrelation function, equation S1 was used

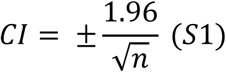

where n is the number of time points[1]. To assess the goodness-of-fitness of the HMM model, the normal pseudo-residuals were used[65], [73]. Assume a set of observations *x*_1_, *x*_2_, …, *x*_*T*_ from a model *X*_*t*_ ∼*F*_*t*_, where *F*_*t*_ is the distribution of random variable *X*_*t*_. The normal pseudo-residuals are defined as:

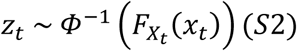

where *ϕ* is the standard normal distribution and *F*_*Xt*_(*x*_*t*_) = Pr (*X*_*t*_ ≤ *x*_*t*_) is the cumulative distribution function or in other words, *x*_*t*_ is transformed to *z*_*t*_. If the fitted model is adequate, then *z*_*t*_ follows a standard normal distribution.

---

**Figure S1.**
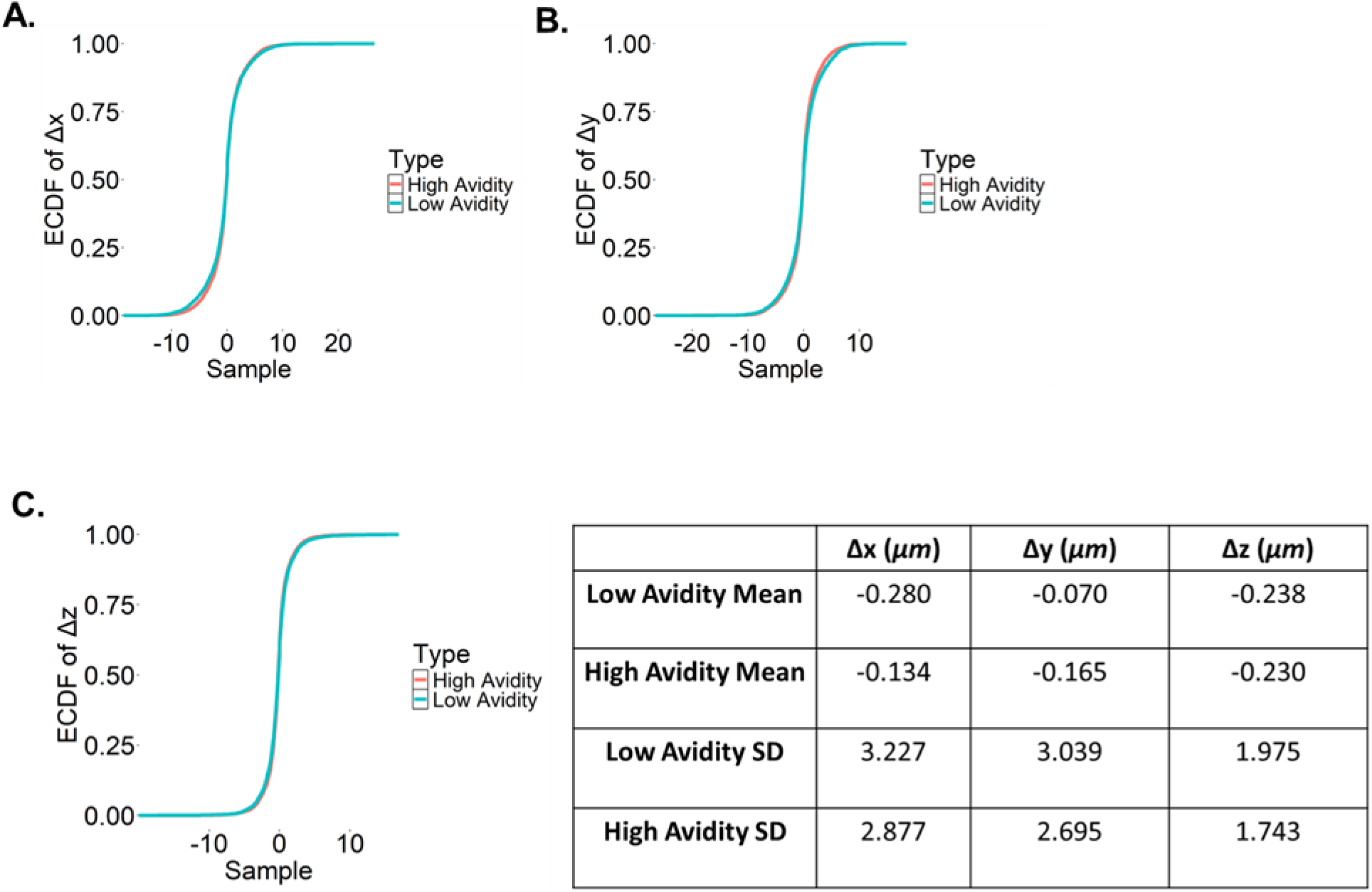
The displacement ECDF of low and high avidity CD8+ T cells on each axis. The p- values are 0.024, 0.0001 and 0.016 for x, y and z axes respectively with a p-value threshold of 0.05, using Kolmogorov-Smirnov test. The table compares the means and the standard deviations of the distributions of the displacements for low and high avidity T cells. High avidity T cells exhibit a lower standard deviation, and it can be attributed to the weaker dissociation rates.

**Figure S2.**
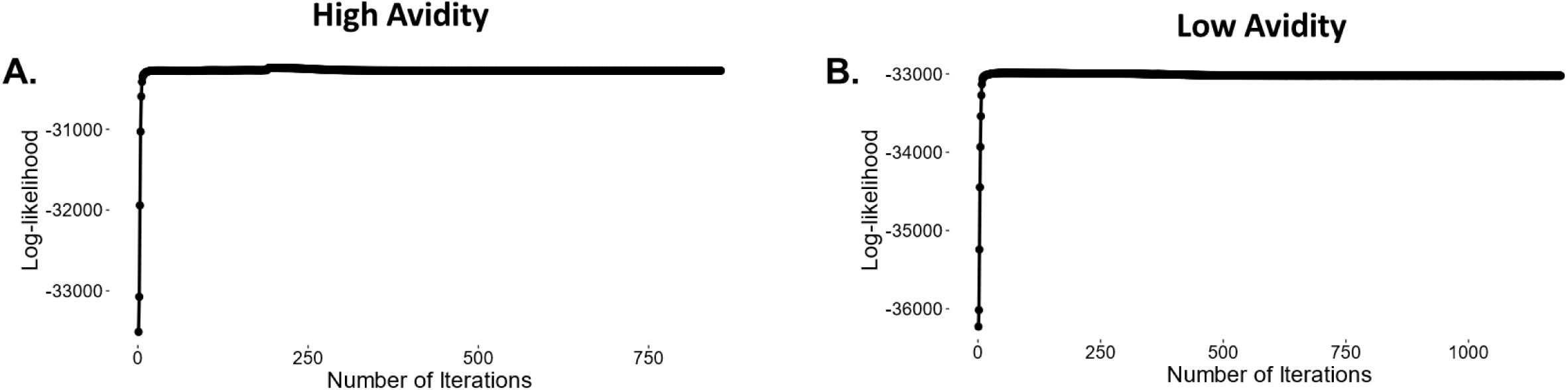
(**A**) Log-likelihood values with respect to the number of iterations till convergence of GMM for high and (**B**) low avidity CD8+ T cells.

**Figure S3.**
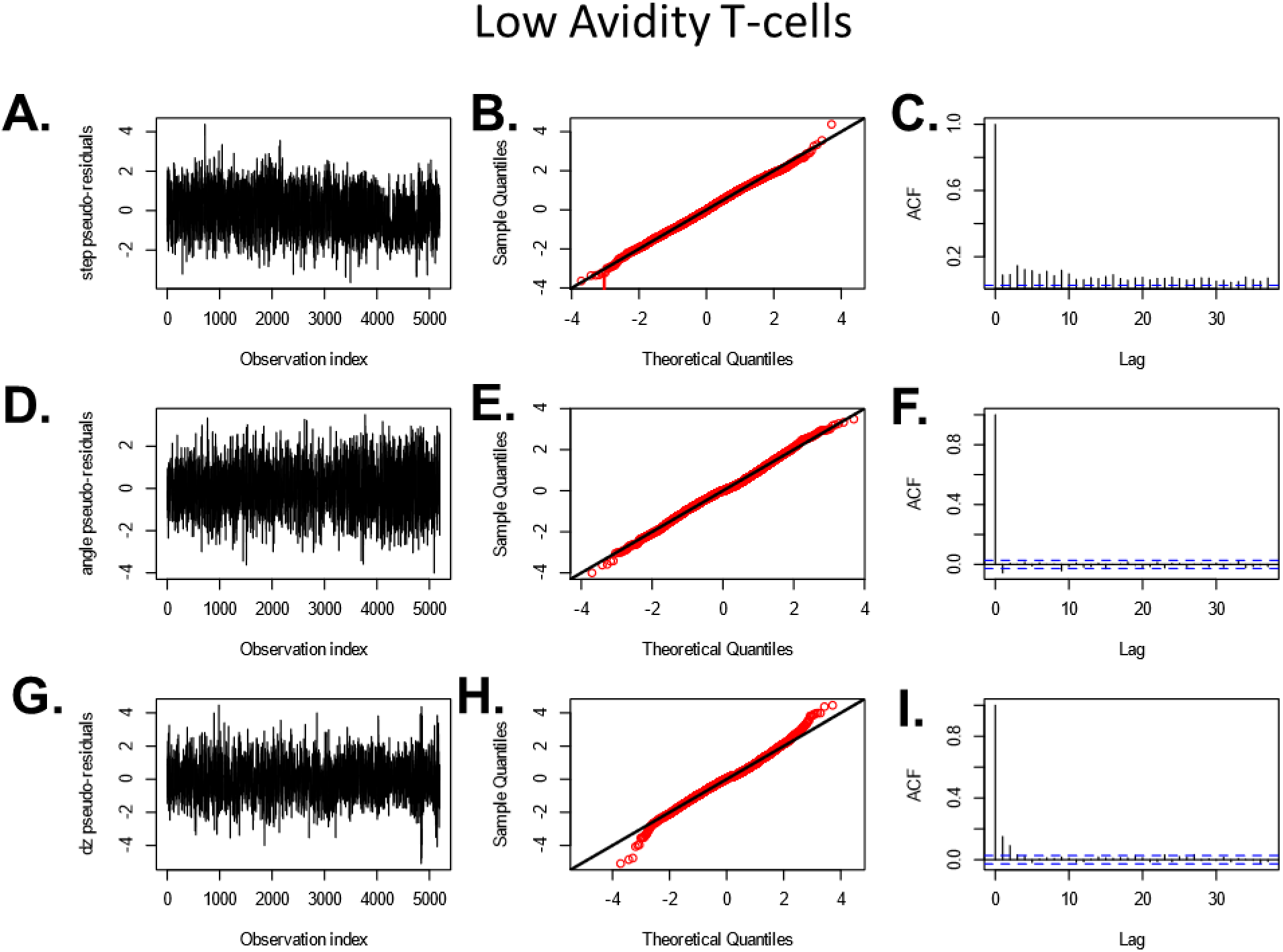
Evaluation of the goodness-of-fit of HMM with pseudo-residuals. (**A**, **D**, **G**) The time series, (**B**, **E**, **H**) the QQ-plots and (**C**, **F**, **I**) the autocorrelation function of the pseudo-residuals for step length, turning angle and Δz respectively for low avidity CD8+ T cells. Deviation from the normal distribution in QQ-plots indicates poor fitting, while high autocorrelation indicates that the model does not capture important factors that influence the migration.

**Figure S4.**
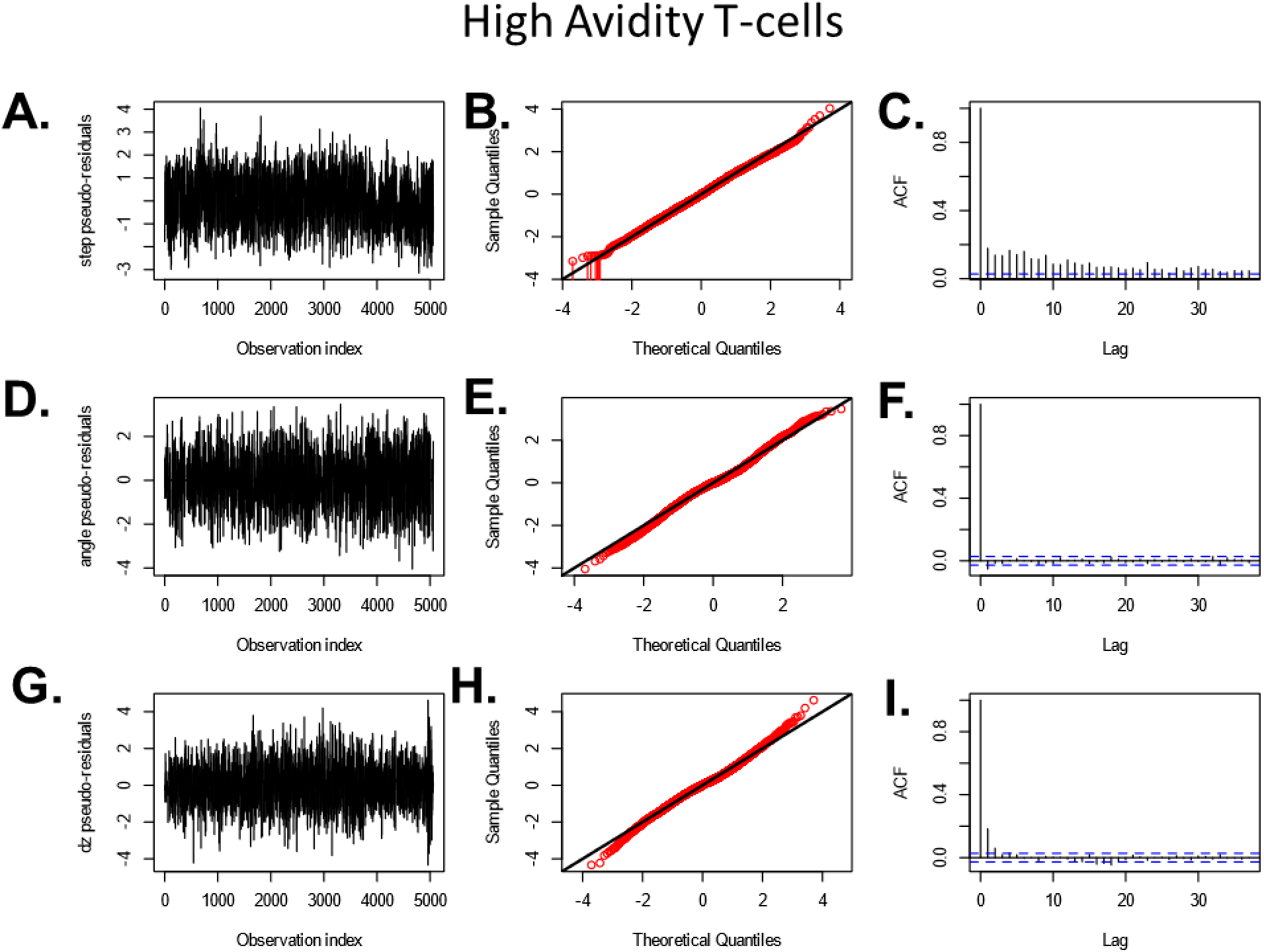
Evaluation of the goodness-of-fit of HMM with pseudo-residuals. (**A**, **D**, **G**) The time series, (**B**, **E**, **H**) the QQ-plots and (**C**, **F**, **I**) the autocorrelation function of the pseudo-residuals for step length, turning angle and Δz respectively for high avidity CD8+ T cells. Deviation from the normal distribution in QQ-plots indicates poor fitting, while high autocorrelation indicates that the model does not capture important factors that influence the migration.

**Figure S5.**
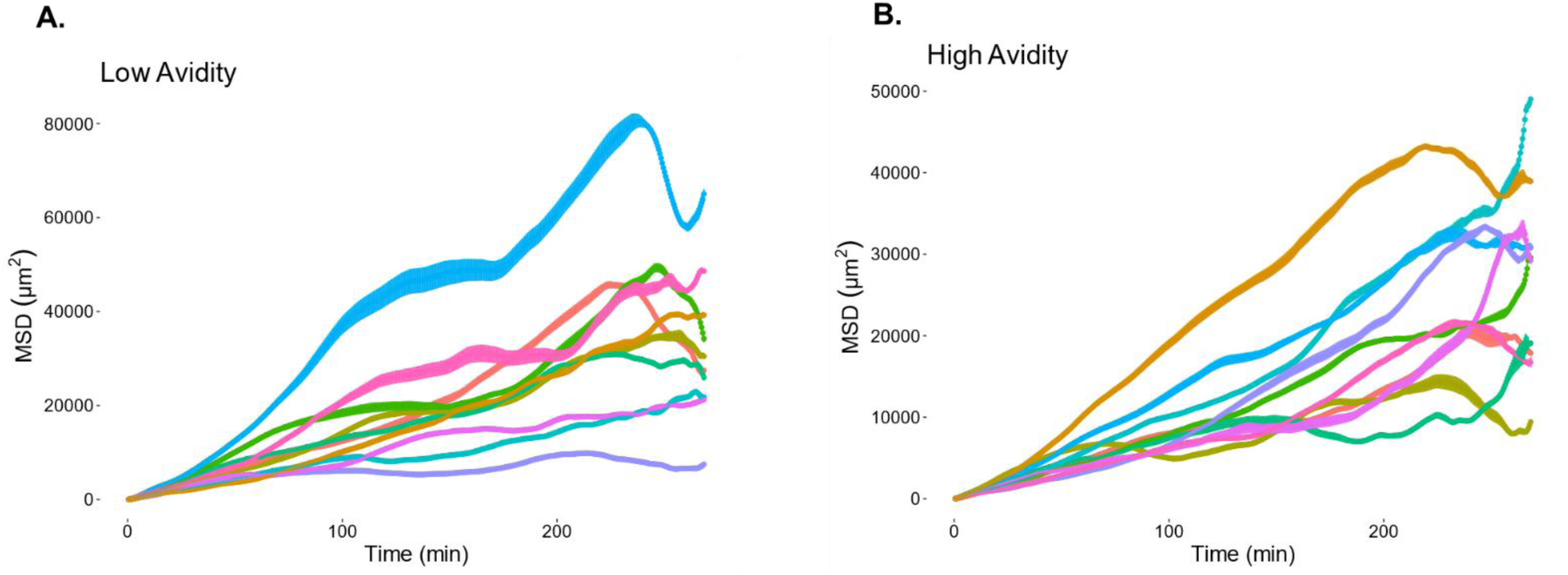
(**A**) Low and (**B**) high avidity time-averaged mean squared displacement plots based on produced simulated data by the Hidden Markov Models.

**Figure S6.**
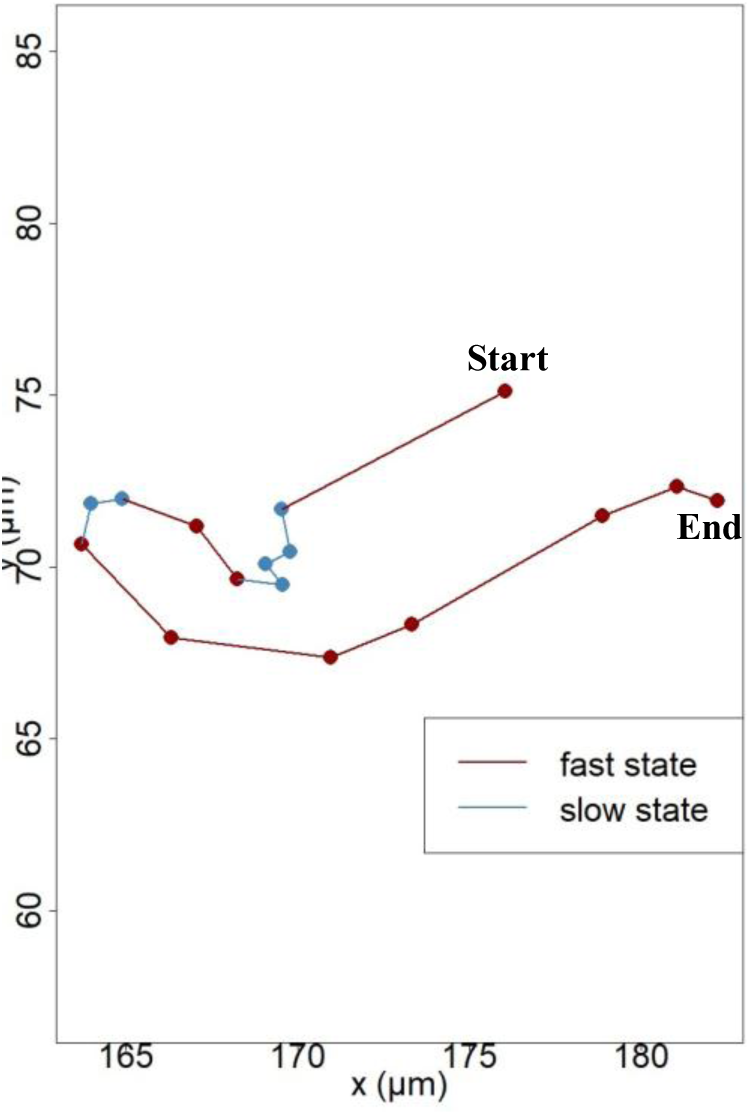
Example of a T cell trajectory and its dynamic switch between the two states.

**Table S1.** Details of the four datasets.

| **Dataset** | **# of Low Avidity T cells** | **# of High Avidity T cells** | **Time interval (sec)** | **Total Duration (sec)** | **Total Duration (min)** |
| --- | --- | --- | --- | --- | --- |
| **1** | 86 | 55 | 31.558 | 1861.92 | 31.032 |
| **2** | 45 | 35 | 33.833 | 1996.15 | 33.26917 |
| **3** | 46 | 100 | 33.833 | 1996.15 | 33.2692 |
| **4** | 48 | 23 | 30.1308 | 1777.72 | 29.6287 |

**Table S2.** Kolmogorov-Smirnov Test p-values for comparing low and high avidity CD8+ T cell distribution of the displacements.

|  | **D** | **p-value** |
| --- | --- | --- |
| **$\Delta x$** | 0.0302 | 0.0238 |
| **$\Delta y$** | 0.0443 | 0.0001 |
| **$\Delta z$** | 0.0316 | 0.01578 |

**Table S3.** Values of the Anderson-Darling Normality Test for the whole CD8+ T cell population, low and high avidity CD8+ T cells separated.

| **Anderson-Darling Normality Test** |  |  |  |
| --- | --- | --- | --- |
|  | **A** | **p-value** | **Axis** |
| **Total** | 224.86 | $2.20 \times 10^{-16}$ | **x** |
| | 205.16 | $2.20 \times 10^{-16}$ | **y** |
| | 294.85 | $2.20 \times 10^{-16}$ | **z** |
| **High Avidity** | 105.13 | $2.20 \times 10^{-16}$ | **x** |
| | 119.44 | $2.20 \times 10^{-16}$ | **y** |
| | 144.19 | $2.20 \times 10^{-16}$ | **z** |
| **Low Avidity** | 117.15 | $2.20 \times 10^{-16}$ | **x** |
| | 86.243 | $2.20 \times 10^{-16}$ | **y** |
| | 149.11 | $2.20 \times 10^{-16}$ | **z** |

**Table S4.** Step distribution parameters for the low avidity T cells in *μm*.

|  | **Slow Moving** | **Fast Moving** |
| --- | --- | --- |
| **Mean** | 1.2687 | 5.0547 |
| **SD** | 0.9599 | 2.9731 |
| **Zero Mass** | $9.14651 \cdot 10^{-17}$ | 0.0007 |

**Table S5.**
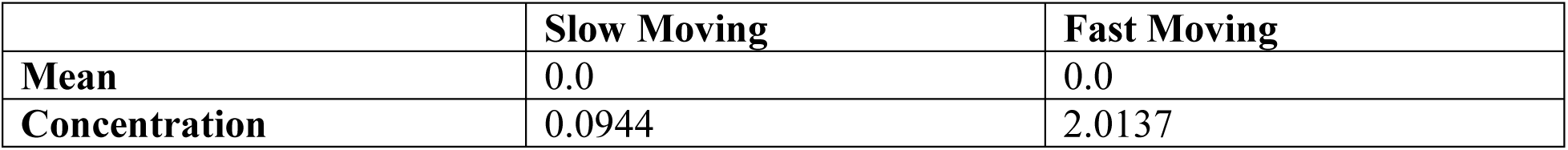
Angle distribution parameters for the low avidity T cells in *radians*.

|  | **Slow Moving** | **Fast Moving** |
| --- | --- | --- |
| **Mean** | 0.0 | 0.0 |
| **Concentration** | 0.0944 | 2.0137 |

**Table S6.**
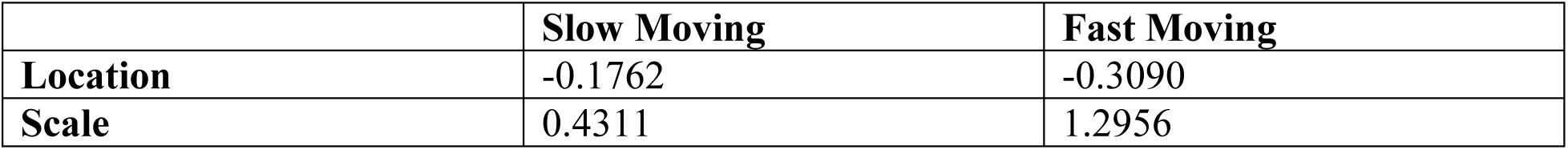
Δz distribution parameters for the low avidity T cells in *μm*.

|  | **Slow Moving** | **Fast Moving** |
| --- | --- | --- |
| **Location** | -0.1762 | -0.3090 |
| **Scale** | 0.4311 | 1.2956 |

**Table S7.**
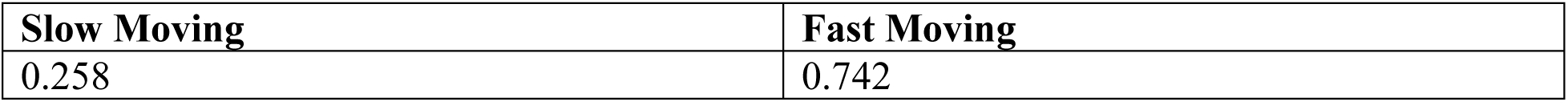
Initial distribution for the low avidity T cells.

**Table S8.** Step distribution parameters for the high avidity T cells in *μm*.

|  | **Slow Moving** | **Fast Moving** |
| --- | --- | --- |
| **Mean** | 1.0801 | 4.3653 |
| **SD** | 0.8089 | 2.7823 |
| **Zero Mass** | 0.0006 | 0.0021 |

**Table S9.** Angle distribution parameters for the high avidity T cells in *radians*.

|  | **Slow Moving** | **Fast Moving** |
| --- | --- | --- |
| **Mean** | 0.0 | 0.0 |
| **Concentration** | 0.1028 | 1.728606 |

**Table S10.** Δz distribution parameters for the high avidity T cells in *μm*.

|  | **Slow Moving** | **Fast Moving** |
| --- | --- | --- |
| **Location** | -0.1998 | -0.2597 |
| **Scale** | 0.3574 | 1.1638 |

**Table S11.** Initial distribution for the high avidity T cells.

| **Slow Moving** | **Fast Moving** |
| --- | --- |
| 0.185 | 0.816 |

## References

[1] X. Trepat, Z. Chen, and K. Jacobson, “Cell Migration,” Compr Physiol, vol. 2, no. 4, pp. 2369–2392, 2012, doi: 10.1002/CPHY.C110012.

[2] W. J. Gao et al., “Macrophage 3D migration: A potential therapeutic target for inflammation and deleterious progression in diseases,” Pharmacol Res, vol. 167, May 2021, doi: 10.1016/J.PHRS.2021.105563.

[3] D. Y. S. Vogel et al., “Macrophages migrate in an activation-dependent manner to chemokines involved in neuroinflammation,” J Neuroinflammation, vol. 11, Feb. 2014, doi: 10.1186/1742-2094-11-23.

[4] P. E. Olofsson et al., “Distinct migration and contact dynamics of resting and IL-2- activated human natural killer cells,” Front Immunol, vol. 5, no. MAR, 2014, doi: 10.3389/FIMMU.2014.00080.

[5] M. F. Krummel, F. Bartumeus, and A. Gérard, “T cell migration, search strategies and mechanisms,” Nature Reviews Immunology 2016 16:*3*, vol. 16, no. 3, pp. 193–201, Feb. 2016, doi: 10.1038/nri.2015.16.

[6] E. D. Tabdanov et al., “Engineering T cells to enhance 3D migration through structurally and mechanically complex tumor microenvironments,” Nature Communications 2021 12:*1*, vol. 12, no. 1, pp. 1–17, May 2021, doi: 10.1038/s41467-021-22985-5.

[7] R. C. Sterner and R. M. Sterner, “CAR-T cell therapy: current limitations and potential strategies,” Blood Cancer Journal 2021 11:*4*, vol. 11, no. 4, pp. 1–11, Apr. 2021, doi: 10.1038/s41408-021-00459-7.

[8] M. H. Kazemi et al., “Tumor-infiltrating lymphocytes for treatment of solid tumors: It takes two to tango?,” Front Immunol, vol. 13, p. 1018962, Oct. 2022, doi: 10.3389/FIMMU.2022.1018962/BIBTEX.

[9] J. Rojas-Quintero et al., “Car T Cells in Solid Tumors: Overcoming Obstacles,” Int J Mol Sci, vol. 25, no. 8, p. 4170, Apr. 2024, doi: 10.3390/IJMS25084170.

[10] S. Abizanda-Campo et al., “Microphysiological systems for solid tumor immunotherapy: opportunities and challenges,” Microsystems & Nanoengineering 2023 9:*1*, vol. 9, no. 1, pp. 1–30, Dec. 2023, doi: 10.1038/s41378-023-00616-x.

[11] I. S. Mauldin et al., “Proliferating CD8+ T cell infiltrates are associated with improved survival in glioblastoma,” Cells, vol. 10, no. 12, p. 3378, Dec. 2021, doi: 10.3390/CELLS10123378/S1.

[12] Y. Yuan et al., “Development and Validation of a CD8+ T Cell Infiltration-Related Signature for Melanoma Patients,” Front Immunol, vol. 12, p. 659444, May 2021, doi: 10.3389/FIMMU.2021.659444/BIBTEX.

[13] L. Chen, Y. Weng, X. Cui, Q. Li, M. Peng, and Q. Song, “Comprehensive analyses of a CD8+ T cell infiltration related gene signature with regard to the prediction of prognosis and immunotherapy response in lung squamous cell carcinoma,” BMC Bioinformatics, vol. 24, no. 1, Dec. 2023, doi: 10.1186/S12859-023-05302-3.

[14] C. H. Polman et al., “A randomized, placebo-controlled trial of natalizumab for relapsing multiple sclerosis,” N Engl J Med, vol. 354, no. 9, pp. 899–910, Mar. 2006, doi: 10.1056/NEJMOA044397.

[15] A. O. Magnan et al., “Assessment of the Th1/Th2 paradigm in whole blood in atopy and asthma. Increased IFN-gamma-producing CD8(+) T cells in asthma,” Am J Respir Crit Care Med, vol. 161, no. 6, pp. 1790–1796, 2000, doi: 10.1164/AJRCCM.161.6.9906130.

[16] S. H. Cho, L. A. Stanciu, S. T. Holgate, and S. L. Johnston, “Increased interleukin-4, interleukin-5, and interferon-gamma in airway CD4+ and CD8+ T cells in atopic asthma,” Am J Respir Crit Care Med, vol. 171, no. 3, pp. 224–230, Feb. 2005, doi: 10.1164/RCCM.200310-1416OC.

[17] O. Lourenço, A. M. Fonseca, and L. Taborda-Barata, “Human CD8+ T Cells in Asthma: Possible Pathways and Roles for NK-Like Subtypes,” Front Immunol, vol. 7, no. DEC, p. 23, 2016, doi: 10.3389/FIMMU.2016.00638.

[18] S. Keshav et al., “A Randomized Controlled Trial of the Efficacy and Safety of CCX282-B, an Orally-Administered Blocker of Chemokine Receptor CCR9, for Patients with Crohn’s Disease,” PLoS One, vol. 8, no. 3, Mar. 2013, doi: 10.1371/JOURNAL.PONE.0060094.

[19] C. R. Mackay, “Moving targets: cell migration inhibitors as new anti-inflammatory therapies,” Nature Immunology 2008 9:*9*, vol. 9, no. 9, pp. 988–998, Aug. 2008, doi: 10.1038/ni.f.210.

[20] E. A. Codling, M. J. Plank, and S. Benhamou, “Random walk models in biology,” Aug. 06, 2008, *Royal Society*. doi: 10.1098/rsif.2008.0014.

[21] Dr. A. Fick, “V. On liquid diffusion,” *The London*, Edinburgh, and Dublin Philosophical Magazine and Journal of Science, vol. 10, no. 63, pp. 30–39, Jul. 1855, doi: 10.1080/14786445508641925.

[22] A. E.-A. der physik and undefined 1906, “Zur theorie der brownschen bewegung,” *scholar.archive.orgA EinsteinAnnalen der physik*, 1906•scholar.archive.org, Accessed: Sep. 18, 2023. [Online]. Available: https://scholar.archive.org/work/ksobd4ymfrf73go3jvjszkjv34/access/ia_file/crossref-pre-1909-scholarly-works/10.1002%252Fandp.19053210303.zip/10.1002%252Fandp.19063240208.pdf

[23] I. M. Sokolov and J. Klafter, “From diffusion to anomalous diffusion: A century after Einstein’s Brownian motion,” Chaos, vol. 15, no. 2, p. 41, Jun. 2005, doi: 10.1063/1.1860472/922577.

[24] H. G. Othmer, S. R. Dunbar, and W. Alt, “Models of dispersal in biological systems,” J Math Biol, vol. 26, no. 3, pp. 263–298, Jun. 1988, doi: 10.1007/BF00277392/METRICS.

[25] G. E. Uhlenbeck and L. S. Ornstein, “On the theory of the Brownian motion,” Physical Review, vol. 36, no. 5, pp. 823–841, 1930, doi: 10.1103/PHYSREV.36.823.

[26] P. H. Wu, A. Giri, S. X. Sun, and D. Wirtz, “Three-dimensional cell migration does not follow a random walk,” Proc Natl Acad Sci U S A, vol. 111, no. 11, pp. 3949–3954, Mar. 2014, doi: 10.1073/PNAS.1318967111/SUPPL_FILE/PNAS.201318967SI.PDF.

[27] P. Dieterich, R. Klages, R. Preuss, and A. Schwab, “Anomalous dynamics of cell migration,” Proc Natl Acad Sci U S A, vol. 105, no. 2, pp. 459–463, Jan. 2008, doi: 10.1073/PNAS.0707603105/ASSET/F2F2348E-1603-4777-A410-613DD4E8A454/ASSETS/GRAPHIC/ZPQ00208-8976-M16.GIF.

[28] H. Gruler, “Locomotion of White Blood Cells: a Biophysical Analysis Langmuir Monolayer View project Clasical Liquid Crystal View project,” 2014. [Online]. Available: https://www.researchgate.net/publication/20389913

[29] Z. Sadjadi, R. Zhao, M. Hoth, B. Qu, and H. Rieger, “Migration of Cytotoxic T Lymphocytes in 3D Collagen Matrices,” Biophys J, vol. 119, no. 11, pp. 2141–2152, Dec. 2020, doi: 10.1016/J.BPJ.2020.10.020.

[30] C. Beauchemin, N. M. Dixit, and A. S. Perelson, “Characterizing T cell movement within lymph nodes in the absence of antigen,” J Immunol, vol. 178, no. 9, pp. 5505–5512, May 2007, doi: 10.4049/JIMMUNOL.178.9.5505.

[31] E. Kepten, A. Weron, G. Sikora, K. Burnecki, and Y. Garini, “Guidelines for the Fitting of Anomalous Diffusion Mean Square Displacement Graphs from Single Particle Tracking Experiments,” PLoS One, vol. 10, no. 2, p. e0117722, Feb. 2015, doi: 10.1371/JOURNAL.PONE.0117722.

[32] A. J. Berglund, “Statistics of camera-based single-particle tracking,” Phys Rev E Stat Nonlin Soft Matter Phys, vol. 82, no. 1, p. 011917, Jul. 2010, doi: 10.1103/PHYSREVE.82.011917/FIGURES/3/MEDIUM.

[33] C. L. Vestergaard, P. C. Blainey, and H. Flyvbjerg, “Optimal estimation of diffusion coefficients from single-particle trajectories,” Phys Rev E Stat Nonlin Soft Matter Phys, vol. 89, no. 2, p. 022726, Feb. 2014, doi: 10.1103/PHYSREVE.89.022726/FIGURES/10/MEDIUM.

[34] H. Qian, M. P. Sheetz, and E. L. Elson, “Single particle tracking. Analysis of diffusion and flow in two-dimensional systems,” Biophys J, vol. 60, no. 4, pp. 910–921, Oct. 1991, doi: 10.1016/S0006-3495(91)82125-7.

[35] G. M. Viswanathan, V. Afanasyev, S. V. Buldyrev, E. J. Murphy, P. A. Prince, and H. E. Stanley, “Lévy flight search patterns of wandering albatrosses,” Nature 1996 381:6581, vol. 381, no. 6581, pp. 413–415, 1996, doi: 10.1038/381413a0.

[36] O. C. Ibe, “Markov Processes for Stochastic Modeling: Second Edition,” Markov Processes for Stochastic Modeling: Second Edition, pp. 1–494, 2013, doi: 10.1016/C2012-0-06106-6.

[37] B. Dybiec, E. Gudowska-Nowak, E. Barkai, and A. A. Dubkov, “Lévy flights versus Lévy walks in bounded domains,” Phys Rev E, vol. 95, no. 5, p. 052102, May 2017, doi: 10.1103/PHYSREVE.95.052102/FIGURES/13/MEDIUM.

[38] M. A. Lomholt, T. Koren, R. Metzler, and J. Klafter, “Lévy strategies in intermittent search processes are advantageous,” Proc Natl Acad Sci U S A, vol. 105, no. 32, pp. 11055– 11059, Aug. 2008, doi: 10.1073/PNAS.0803117105/ASSET/4D7D9440-8D97-49BF-9BF4-829DD681FBB7/ASSETS/GRAPHIC/ZPQ999083502CL004.JPEG.

[39] A. M. Reynolds, “Lévy flight movement patterns in marine predators may derive from turbulence cues,” Proc Math Phys Eng Sci, vol. 470, no. 2171, Nov. 2014, doi: 10.1098/RSPA.2014.0408.

[40] M. S. Abe, “Functional advantages of Lévy walks emerging near a critical point,” Proc Natl Acad Sci U S A, vol. 117, no. 39, pp. 24336–24344, Sep. 2020, doi: 10.1073/PNAS.2001548117/SUPPL_FILE/PNAS.2001548117.SAPP.PDF.

[41] G. Ariel, A. Rabani, S. Benisty, J. D. Partridge, R. M. Harshey, and A. Be’Er, “Swarming bacteria migrate by Lévy Walk,” Nature Communications 2015 6:*1*, vol. 6, no. 1, pp. 1–6, Sep. 2015, doi: 10.1038/ncomms9396.

[42] I. Rhee, M. Shin, S. Hong, K. Lee, and S. Chong, “On the Levy-Walk Nature of Human Mobility,” pp. 924–932, Jun. 2008, doi: 10.1109/INFOCOM.2008.145.

[43] T. H. Harris et al., “Generalized Lévy walks and the role of chemokines in migration of effector CD8+ T cells,” Nature, vol. 486, no. 7404, p. 545, Jun. 2012, doi: 10.1038/NATURE11098.

[44] A. James, M. J. Plank, and A. M. Edwards, “Assessing Lévy walks as models of animal foraging,” J R Soc Interface, vol. 8, no. 62, pp. 1233–1247, Sep. 2011, doi: 10.1098/RSIF.2011.0200.

[45] A. M. Edwards, M. P. Freeman, G. A. Breed, and I. D. Jonsen, “Incorrect Likelihood Methods Were Used to Infer Scaling Laws of Marine Predator Search Behaviour,” PLoS One, vol. 7, no. 10, p. e45174, Oct. 2012, doi: 10.1371/JOURNAL.PONE.0045174.

[46] A. M. Edwards, “Overturning conclusions of Lévy flight movement patterns by fishing boats and foraging animals,” Ecology, vol. 92, no. 6, pp. 1247–1257, Jun. 2011, doi: 10.1890/10-1182.1.

[47] E. J. Banigan, T. H. Harris, D. A. Christian, C. A. Hunter, and A. J. Liu, “Heterogeneous CD8+ T Cell Migration in the Lymph Node in the Absence of Inflammation Revealed by Quantitative Migration Analysis,” PLoS Comput Biol, vol. 11, no. 2, p. e1004058, 2015, doi: 10.1371/JOURNAL.PCBI.1004058.

[48] E. R. Jerison and S. R. Quake, “Heterogeneous T cell motility behaviors emerge from a coupling between speed and turning in vivo,” Elife, vol. 9, pp. 1–26, May 2020, doi: 10.7554/ELIFE.53933.

[49] B.-J. Yoon, “Hidden Markov Models and their Applications in Biological Sequence Analysis,” Curr Genomics, vol. 10, no. 6, p. 402, Sep. 2009, doi: 10.2174/138920209789177575.

[50] L. R. Rabiner, “A Tutorial on Hidden Markov Models and Selected Applications in Speech Recognition,” Proceedings of the IEEE, vol. 77, no. 2, pp. 257–286, 1989, doi: 10.1109/5.18626.

[51] J. Degerman et al., “An automatic system for in vitro cell migration studies,” J Microsc, vol. 233, no. 1, pp. 178–191, 2009, doi: 10.1111/J.1365-2818.2008.03108.X.

[52] F. Mohammadi et al., “A lineage tree-based hidden Markov model quantifies cellular heterogeneity and plasticity,” Communications Biology 2022 5:*1*, vol. 5, no. 1, pp. 1–14, Nov. 2022, doi: 10.1038/s42003-022-04208-9.

[53] M. Held et al., “CellCognition: time-resolved phenotype annotation in high-throughput live cell imaging,” Nature Methods 2010 7:*9*, vol. 7, no. 9, pp. 747–754, Aug. 2010, doi: 10.1038/nmeth.1486.

[54] S. Gordonov, M. K. Hwang, A. Wells, F. B. Gertler, D. A. Lauffenburger, and M. Bathe, “Time series modeling of live-cell shape dynamics for image-based phenotypic profiling,”Integrative Biology, vol. 8, no. 1, pp. 73–90, Jan. 2016, doi: 10.1039/C5IB00283D.

[55] E. Torkashvand, “Modeling three-dimensional T-cell motility using clustering and hidden Markov models,” Stat Methods Med Res, vol. 32, no. 7, Jul. 2023, doi: 10.1177/09622802231172041.

[56] C. G. Tucker et al., “Adoptive T Cell Therapy with IL-12–Preconditioned Low-Avidity T Cells Prevents Exhaustion and Results in Enhanced T Cell Activation, Enhanced Tumor Clearance, and Decreased Risk for Autoimmunity,” The Journal of Immunology, vol. 205, no. 5, pp. 1449–1460, Sep. 2020, doi: 10.4049/JIMMUNOL.2000007.

[57] D. I. Shreiber, V. H. Barocas, and R. T. Tranquillo, “Temporal Variations in Cell Migration and Traction during Fibroblast-Mediated Gel Compaction,” Biophys J, vol. 84, no. 6, p. 4102, Jun. 2003, doi: 10.1016/S0006-3495(03)75135-2.

[58] D. J. Odde,’, E. M. Tanaka, S. S. Hawkins,’, and H. M. Buettner’zt, “Stochastic Dynamics of the Nerve Growth Cone and Its Microtubules During Neurite Outgrowth”, doi: 10.1002/(SICI)1097-0290(19960520)50:4.

[59] “Pattern Recognition and Machine Learning,” *Pattern Recognition and Machine Learning*, Dec. 2006, doi: 10.1007/978-0-387-45528-0.

[60] C. Shi, B. Wei, S. Wei, W. Wang, H. Liu, and J. Liu, “A quantitative discriminant method of elbow point for the optimal number of clusters in clustering algorithm,” EURASIP J Wirel Commun Netw, vol. 2021, no. 1, pp. 1–16, Dec. 2021, doi: 10.1186/S13638-021-01910-W/FIGURES/6.

[61] B. T. McClintock and T. Michelot, “momentuHMM: R package for generalized hidden Markov models of animal movement,” Methods Ecol Evol, vol. 9, no. 6, pp. 1518–1530, Jun. 2018, doi: 10.1111/2041-210X.12995.

[62] T. Michelot, R. Langrock, and T. A. Patterson, “moveHMM: an R package for the statistical modelling of animal movement data using hidden Markov models,” Methods Ecol Evol, vol. 7, no. 11, pp. 1308–1315, Nov. 2016, doi: 10.1111/2041-210X.12578.

[63] T. Michelot, R. Langrock, and T. A. Patterson, “moveHMM: an R package for the statistical modelling of animal movement data using hidden Markov models,” Methods Ecol Evol, vol. 7, no. 11, pp. 1308–1315, Nov. 2016, doi: 10.1111/2041-210X.12578.

[64] “Numerical Maximisation of Likelihood: A Neglected Alternative to EM? on JSTOR.” Accessed: Dec. 17, 2023. [Online]. Available: https://www.jstor.org/stable/43299760

[65] W. Zucchini, I. L. MacDonald, and R. Langrock, Hidden Markov models for time series : an introduction using R. Accessed: Dec. 17, 2023. [Online]. Available: https://www.routledge.com/Hidden-Markov-Models-for-Time-Series-An-Introduction-Using-R-Second-Edition/Zucchini-MacDonald-Langrock/p/book/9781032179490

[66] P. Mrass, J. Petravic, M. P. Davenport, and W. Weninger, “Cell-autonomous and environmental contributions to the interstitial migration of T cells,” Semin Immunopathol, vol. 32, no. 3, pp. 257–274, Sep. 2010, doi: 10.1007/S00281-010-0212-1/TABLES/5.

[67] C. Beauchemin, N. M. Dixit, and A. S. Perelson, “Characterizing T Cell Movement within Lymph Nodes in the Absence of Antigen,” The Journal of Immunology, vol. 178, no. 9, pp. 5505–5512, May 2007, doi: 10.4049/JIMMUNOL.178.9.5505.

[68] S. P. Preston, S. L. Waters, O. E. Jensen, P. R. Heaton, and D. I. Pritchard, “T-cell motility in the early stages of the immune response modeled as a random walk amongst targets,” Phys Rev E Stat Nonlin Soft Matter Phys, vol. 74, no. 1 Pt 1, 2006, doi: 10.1103/PHYSREVE.74.011910.

[69] M. F. Krummel, F. Bartumeus, and A. Gérard, “T cell migration, search strategies and mechanisms,” Nature Reviews Immunology 2016 16:*3*, vol. 16, no. 3, pp. 193–201, Feb. 2016, doi: 10.1038/nri.2015.16.

[70] C. M. Witt, S. Raychaudhuri, B. Schaefer, A. K. Chakraborty, and E. A. Robey, “Directed migration of positively selected thymocytes visualized in real time,” PLoS Biol, vol. 3, no. 6, pp. 1062–1069, 2005, doi: 10.1371/JOURNAL.PBIO.0030160.

[71] A. James, M. J. Plank, and A. M. Edwards, “Assessing Lévy walks as models of animal foraging,” J R Soc Interface, vol. 8, no. 62, pp. 1233–1247, Sep. 2011, doi: 10.1098/RSIF.2011.0200.

[72] H. Sakoe and S. Chiba, “Dynamic Programming Algorithm Optimization for Spoken Word Recognition,” IEEE Trans Acoust, vol. 26, no. 1, pp. 43–49, 1978, doi: 10.1109/TASSP.1978.1163055.

[73] A. Stadie, “Überprüfung stochastischer Modelle mit Pseudo-Residuen,” Feb. 2003, doi: 10.53846/GOEDISS-3671.

## References

[1] R. J. Hyndman and G. Athanasopoulos, “Forecasting: Principles and Practice,” 2018, *OTexts*. Accessed: Oct. 27, 2024. [Online]. Available: https://research.monash.edu/en/publications/forecasting-principles-and-practice-2

[2] W. Zucchini, I. L. MacDonald, and R. Langrock, Hidden Markov models for time series : an introduction using R. Accessed: Dec. 17, 2023. [Online]. Available: https://www.routledge.com/Hidden-Markov-Models-for-Time-Series-An-Introduction-Using-R-Second-Edition/Zucchini-MacDonald-Langrock/p/book/9781032179490

[3] A. Stadie, “Überprüfung stochastischer Modelle mit Pseudo-Residuen,” Feb. 2003, doi: 10.53846/GOEDISS-3671.

